# One-click image reconstruction in single-molecule localization microscopy via deep learning

**DOI:** 10.1101/2025.04.13.648574

**Authors:** Alon Saguy, Dafei Xiao, Kaarjel K. Narayanasamy, Yuya Nakatani, Anna-Karin Gustavsson, Mike Heilemann, Yoav Shechtman

**Affiliations:** Faculty of Biomedical Engineering, Technion – Israel Institute of Technology, Haifa, Israel; EMBL Imaging Centre, European Molecular Biology Laboratory, Heidelberg, Germany; Institute of Physical and Theoretical Chemistry, Goethe-University Frankfurt, Germany; Faculty of Electrical and Computer Engineering Technion – Israel Institute of Technology Haifa, Israel; Walker Department of Mechanical Engineering, University of Texas at Austin, Austin, TX, USA; Russell Berrie Nanotechnology Institute, Technion–Israel Institute of Technology, Haifa, Israel; Department of Chemistry, Rice University, Houston, TX, USA; Department of BioSciences, Rice University, Houston, TX, USA; Department of Electrical and Computer Engineering, Rice University, Houston, TX, USA; Smalley-Curl Institute, Rice University, Houston, TX, USA; Center for Nanoscale Imaging Sciences, Rice University, Houston, TX, USA; Department of Cancer Biology, University of Texas MD Anderson Cancer Center, Houston, TX, USA

## Abstract

Deep neural networks have led to significant advancements in microscopy image generation and analysis. In single-molecule localization based super-resolution microscopy, neural networks are capable of predicting fluorophore positions from high-density emitter data, thus reducing acquisition time, and increasing imaging throughput. However, neural network-based solutions in localization microscopy require intensive human intervention and computation expertise to address the compromise between model performance and its generalization. For example, researchers manually tune parameters to generate training images that are similar to their experimental data; thus, for every change in the experimental conditions, a new training set should be manually tuned, and a new model should be trained. Here, we introduce AutoDS and AutoDS3D, two software programs for reconstruction of single-molecule super-resolution microscopy data that are based on Deep-STORM and DeepSTORM3D, that significantly reduce human intervention from the analysis process by automatically extracting the experimental parameters from the imaging raw data. In the 2D case, AutoDS selects the optimal model for the analysis out of a set of pre-trained models, hence, completely removing user supervision from the process. In the 3D case, we improve the computation efficiency of DeepSTORM3D and integrate the lengthy workflow into a graphic user interface that enables image reconstruction with a single click. Ultimately, we demonstrate superior performance of both pipelines compared to Deep-STORM and DeepSTORM3D for single-molecule imaging data of complex biological samples, while significantly reducing the manual labor and computation time.

## Introduction

Super-resolution microscopy enhances the resolving capability of optical imaging to interrogate length scales below the diffraction limit of light. In single-molecule localization microscopy (SMLM) methods, such as direct stochastic optical reconstruction microscopy ((*d*)STORM)^1,2^, DNA-points accumulation for imaging in nanoscale topography (DNA-PAINT)^3^, and (fluorescence) photoactivated localization microscopy ((f)PALM)^4,5^, the localization of non-overlapping fluorophore signals yields an improvement in resolution of an order of magnitude or more^6^. Furthermore, SMLM can be extended to 3D imaging by employing point spread function (PSF) engineering^7,8^, where the optical system is altered to encode the axial positions of fluorophores in the PSF shape. Since SMLM relies on accumulating isolated emitter signals to generate a super-resolved image, these methods naturally result in long imaging times. Consequentially, the throughput of SMLM methods is limited, and live-cell imaging is restricted to slow biological dynamics.

The integration of neural networks (NNs) into SMLM reduces the acquisition time by enabling the prediction of super-resolved images from high-emitter density imaging data^9–14^. Typically, the training data used to train these networks is based on images that highly resemble the experimental data; the level of similarity between the training data and the experimental data strongly affects performance. One strategy to prepare such training data is to generate a large and diverse simulated dataset^10–12^, which depends mainly on selecting optimal simulation parameters, such as noise levels, emitter density, as well as the encoding PSF^15^, in the 3D case. Alternatively, the application of exchangeable fluorophore labels^16^ in an SMLM experiment allows one to generate experimental data for model training, as demonstrated using Deep-STORM in combination with DNA-PAINT^17^ and live-cell HaloTag-PAINT^18^.

In 2D SMLM, current NNs that predict super-resolved images from high-emitter density images face a common challenge: dealing with heterogeneous data, including varying emitter densities, signal-to-noise ratio (SNR) conditions, and background levels within a single dataset, or even within a single frame; the performance of a single trained model applied across a heterogeneous dataset is limited. Furthermore, models with narrow applicability across datasets are particularly limited for image restoration packages that are sensitive to variation in noise profiles, and model retraining is required with even relatively minor changes to acquisition parameters^19^. This bottleneck introduces a notable challenge in balancing the generalizability of the network across different sample conditions and its overall performance, a challenge also known as the bias-variance trade-off. Interestingly, apart from increasing the training data variability for single models^10^, no existing learning-based localization algorithm has adequately addressed the issue of dataset heterogeneity.

In the context of 3D SMLM by PSF engineering, although the heterogeneity issue exists there as well^20,21^, a more limiting problem is that the whole analysis pipeline, from pre-processing to post-processing is extremely manual labor demanding. The process typically includes: accurate PSF characterization, image pre-processing, training data generation, network training, inference, and post-processing, involving multiple programming languages and software. Furthermore, many parameters in this process need to be properly and manually tuned to guarantee the model’s performance, which requires user expertise. Overall, this process is time-consuming and challenging for end users.

In this work, we demonstrate several contributions to the automation of deep learning-based 2D and 3D localization microscopy. In the 2D case, 1) we develop a fully-automated algorithm for experimental parameter extraction that alleviates the need for manual tuning of SMLM training data; 2) we provide a set of pre-trained Deep-STORM models that covers a wide range of experimental conditions and completely eliminates the need for user intervention during inference; 3) we show that feeding Deep-STORM with patches of the field-of-view and analyzing them separately leads to improved performance compared to processing of entire frames. Our results show significant improvement in predicting super-resolution images from heterogenous high density SMLM datasets of complex biological samples compared to using a single, frame-wide model. In the 3D case, we automate the standard image processing pipeline of PSF engineering-based SMLM and develop AutoDS3D, a one-click graphic user interface (GUI) built on the DeepSTORM3D (DS3D) framework^11^. AutoDS3D achieves reconstruction quality comparable to DeepSTORM3D across various PSF types while operating faster and fully automatically. Overall, these contributions extend the previous Deep-STORM framework; hence, we refer to our method here as AutoDS and AutoDS3D.

Notably, our contribution extends beyond algorithm performance and includes the capability of using AutoDS/3D as a first-of-its-kind one-click plug-and-play tool for non-expert users, significantly removing human decision-making from the loop, and in the case of 2D imaging, completely alleviating the need for post-experiment model training. This contributes to the democratization of state-of-the-art deep learning tools by significantly reducing the human-supervised model-training step and improving prediction quality.

## Results

### AutoDS

Our analysis scheme starts with the division of each input video into multiple patches (Figure 1). The choice of patch size is important: undersized patches lead to suboptimal decisions based on local properties, e.g., either zero emitter density in the absence of emitters in the patch, or very high emitter density when only a single emitter is present. On the other hand, the use of oversized patches ultimately converges to the global parameter-based network selection. Another important trade-off governed by the choice of patch size is between runtime and the model optimization granularity. Although smaller patches are processed faster than large patches by convolutional NNs, decreasing the patch size leads to more forward passes through the model and longer runtimes eventually. In this study, we selected patch dimensions of 14.9 *µm* x 14.9 *µm* (64 x 64 pixels), which we found to be a good compromise between the aforementioned considerations. Nevertheless, our empirical experience showed that this pipeline is robust to patch lengths in the range of 2 – 15 *µm*.

**Figure 1:**
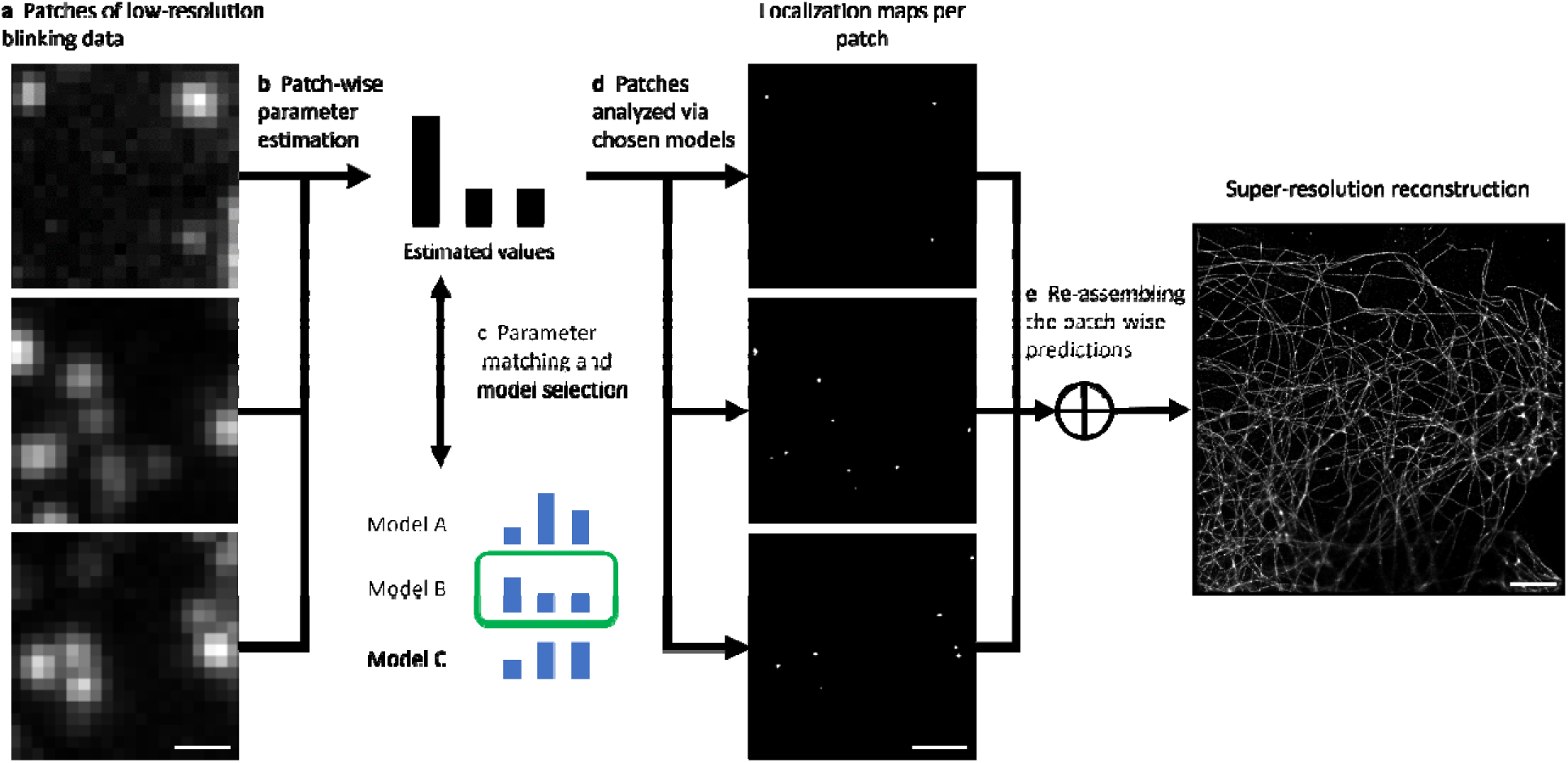
AutoDS workflow: **a** dividing the experimental SMLM video into multiple patches; **b** extracting the emitter density and the SNR parameters from each patch, and comparing the estimated patch parameters to the parameters used for the pre-trained models; **c** choosing the best fitting model to analyze each patch (green box marks the chosen model); **d** analyzing the patches using the selected pre-trained Deep-STORM model and generating the localization maps; **e** reassembling the patches according to their original position in the FOV to obtain a super-resolution reconstruction (bottom: diffraction limited image). Scale bars are 800 nm (**a, b**) and 5 µm (**e**).

Crucially, we have made AutoDS independent of wavelength, numerical aperture (NA), and camera pixel size, by allowing users to specify these parameters during model training and during inference. This allows us to use pre-trained models, by automatically down/up-scaling the experimental images during inference.

Subsequently, we employ an automated parameter-extraction algorithm to classify the patches based on emitter density and the mean and standard deviation of the following properties (see Methods section for more details on parameter extraction): background, noise, PSF photon intensity, and PSF dimensions (see Supplementary figure S1 for visualization of the estimation performance of the model). Based on these classifications, the appropriate model from a set of four pre-trained Deep-STORM models is chosen for processing each patch. Finally, each patch is analyzed by the corresponding model, and the resulting localization maps are integrated into the full field-of-view (FOV). The whole pipeline requires no decision-making by the user, making it user-friendly and suitable for non-experts.

We applied AutoDS to high-density DNA-PAINT data recorded from rat brain tissue labeled for two targets, TOM20 and α-tubulin^17^ (Figure 2a,b). Per target, imager strand concentrations were tuned to obtain high-emitter densities. Our automated pipeline estimated that the emitter densities were in the range of [0.07, 2.19] emitters per µm^2^ (see Supplementary Table S1 for estimated parameter statistics). The targets were recorded for 400 frames and at single frame acquisition time of 150 ms, corresponding to a 1-minute total imaging time. From these datasets, super-resolved images were predicted with AutoDS. Moreover, a ground truth dataset was acquired for the same samples with low emitter density at a total imaging time of 25 minutes. The ground truth images were reconstructed using an SMLM fitting software^22^.

**Figure 2:**
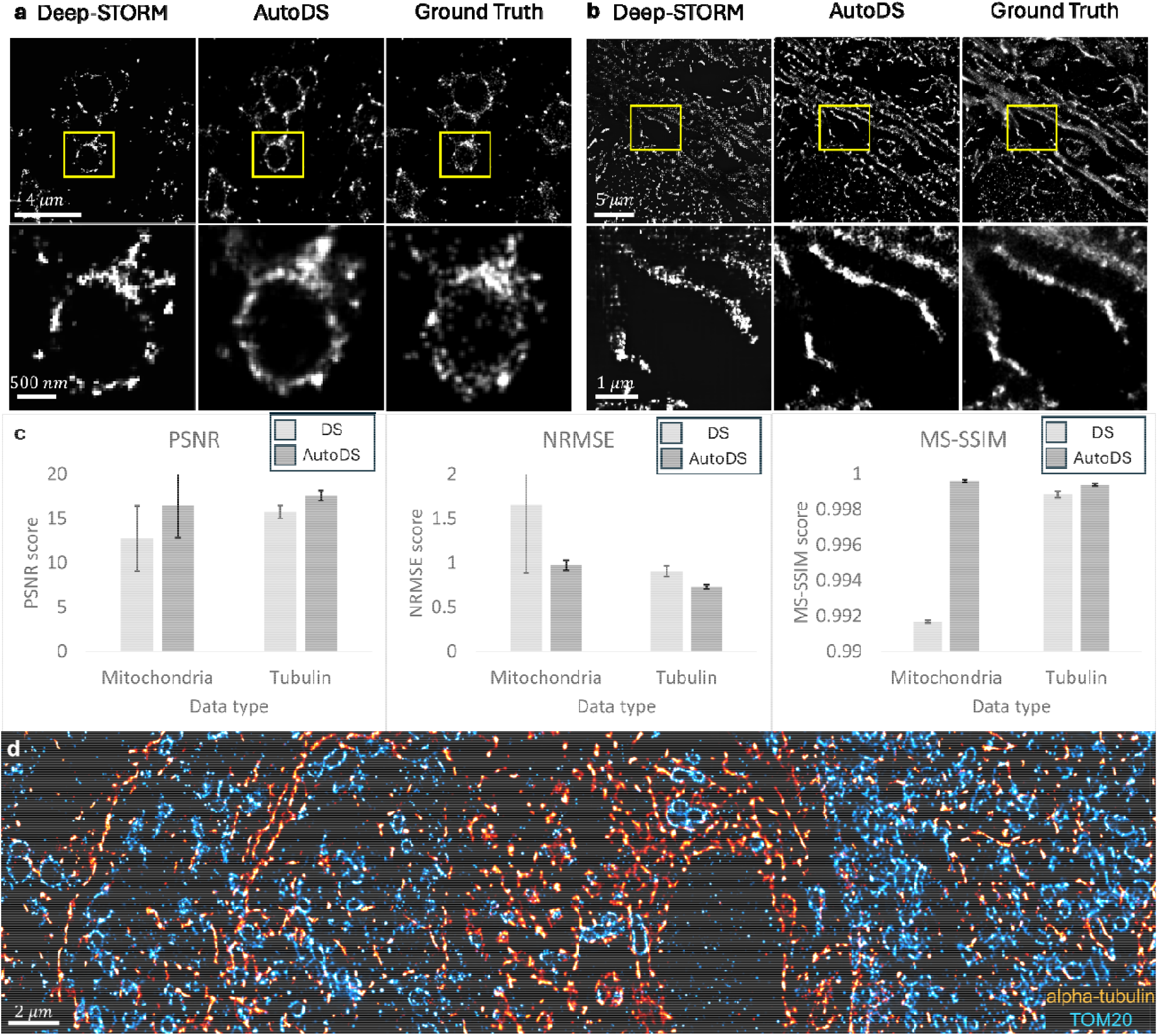
Reconstruction quality of AutoDS. Comparison between Deep-STORM (left), AutoDS (middle), and ground truth (right) reconstructions of **a** TOM20 and **b** α-tubulin. We selected the most frequentl used Deep-STORM model for the comparison to AutoDS. **c** a comparison of the mean PSNR, NRMSE, and MS-SSIM (error bars = standard deviation values) across n=5 different experiments obtained by Deep-STORM reconstructions vs AutoDS reconstructions. **d** Two-color AutoDS reconstruction of a rat brain tissue section labeled for alpha-tubulin (Red) and TOM20 (Blue) and imaged with DNA-PAINT.

We compared the patchwise reconstructions of AutoDS to the reconstruction of a single Deep-STORM model that processed the entire field-of-view. Visual inspection of the predicted images generated by AutoDS versus the ground truth images (Figure 2a,b) showed faithful reconstruction in structure and density of targets. To quantitatively assess the performance of AutoDS we measured the similarity between the reconstructed images and the ground truth images using the peak signal to noise ratio (PSNR), the normalized root mean squared error (NRMSE), and the multi-scale structural similarity index (MS-SSIM)^23,24^.

Qualitatively, the AutoDS reconstructions show superior visual quality compared to the single-model approach, with a higher number of predicted emitters, resulting in more continuous structural features in some regions. Quantitatively, all metrics showed that AutoDS outperforms the single-model approach (see Table 1), supporting the visual observations (Figure 2c). Nevertheless, the reconstruction quality remained similar in most of the field-of-view, mainly because the analyzed samples possessed the same emitter density and SNR throughout most of the field-of-view. In other words, the highest improvement of our method is shown in regions where the emitter densities and SNR deviate from their mean values (see supplementary figure S2).

**Table 1:** Quantitative comparison between DeepSTORM and AutoDS reconstructions on the mitochondria and tubulin datasets. The mean PSNR, NRMSE, and MS-SSIM are reported with ± one standard deviation over n=5 different fields of view.

| Reconstruction algorithm | Sample type | PSNR | NRMSE | MS-SSIM |
| --- | --- | --- | --- | --- |
| Deep-STORM | Mitochondria | $12.74 \pm 3.65$ | $1.65 \pm 0.76$ | $0.991 \pm 6.8 \cdot 10^{-5}$ |
| AutoDS | Mitochondria | $16.44 \pm 0.56$ | $0.97 \pm 0.05$ | $0.999 \pm 7.5 \cdot 10^{-5}$ |
| Deep-STORM | Tubulin | $15.71 \pm 0.71$ | $0.9 \pm 0.06$ | $0.998 \pm 1.7 \cdot 10^{-4}$ |
| AutoDS | Tubulin | $17.59 \pm 0.56$ | $0.73 \pm 0.02$ | $0.999 \pm 7.9 \cdot 10^{-5}$ |

We examined the distribution of the selected pre-trained models for both the TOM20 and α-tubulin experiments and found that the model selection distribution was spread across multiple models. This finding shows the potential contribution of our pipeline for the analysis of experimental data with high variability in sample properties like emitter density and SNR. Moreover, having a set of pre-trained models suitable for multiple experimental conditions is highly beneficial to remove user intervention from the analysis process. An application that demands such a flexibility is multi-target imaging (Figure 2d), where AutoDS can serve as a digital add-on to a microscope setup and can enable straight-forward multiplexing^25^.

A major implication of our automated framework is speeding-up of SMLM analysis, towards potentially real-time SMLM. This is achievable due to the complete elimination of any post-experiment training requirements. To quantify the speed enhancement, we analyzed a typical SMLM video containing 400 high-emitter density frames of size = 512 x 512 pixels, which contained sufficient information for a complete structural reconstruction. For this dataset the model selection took less than a minute, and the inference and reassembly of the reconstructed image from the patches took another 5 minutes. This is an order of magnitude faster compared to a neural network that requires training post-experiment^9^ (typically ∼1 hour per model training).

### AutoDS3D

AutoDS3D is a web application that runs on either a local computer or a remote server. It takes optical parameters, a calibration z-stack, and raw images from SMLM data set as input, and generates a localization list with a single click or step-by-step interactions (Supplementary figure S3). To enhance computation efficiency, AutoDS3D employs Gaussian representation of emitters in space. Additionally, like AutoDS, it automates the image analysis workflow, eliminating the need for technically demanding manual labor.

DS3D^11^ discretizes 3D object space into voxels, assigning a value of 1 to voxels occupied by an emitter and 0 elsewhere. This binary representation limits precision to the voxel size, which means that high resolution demand large matrices with small voxels, which significantly increases computational complexity. AutoDS3D addresses this limitation by representing emitters using 3D Gaussians. The center of mass of such a Gaussian defines emitter location, enabling the use of larger voxels while maintaining high localization precision and improving computational efficiency.

For automation, we integrate: a phase retrieval algorithm for PSF characterization^15,26^, image pre-processing for background removal, SNR characterization, training data generation, network training, inference test and final inference on all the images. Intermediate results are generated during the execution to provide feedback (Supplementary figure S4). This step completely alleviates the extremely time- and expertise-demanding need for user parameter selection (e.g. SNR, emitter density) by eyeballing and trial and error.

We first compared the performance of 1) AutoDS3D; 2) DS3D with; and 3) without 4x up-sampling for lateral voxel size control. These three models are trained under controlled conditions including the PSF model, the number of training images, and the level of noise and photon count. We then performed 1000 inference runs with each trained model to assess localization performance, on simulated images of randomly located emitters in 3D.

The evaluation metrics include: duration of training data generation, training duration, inference duration, post-processing duration, Jaccard index, and lateral and axial root mean square error (RMSE) between the localizations and simulated ground truth locations. This comparison is conducted under both high and low SNR conditions, as illustrated in the radar plots of Figure 3. In high SNR condition (Figure 3a), AutoDS3D achieves the best computation efficiency and Jaccard index among the three models while maintaining localization accuracy comparable to DS3D. Although DS3D without up-sampling reduces operation time, it compromises lateral RMSE due to the coarse lateral voxel size. Under low SNR condition (Figure 3b), AutoDS3D continues to lead in computational efficiency and Jaccard index, while localization RMSE becomes comparable across all models, likely due to the dominant influence of noise on localization precision.

**Figure 3:**
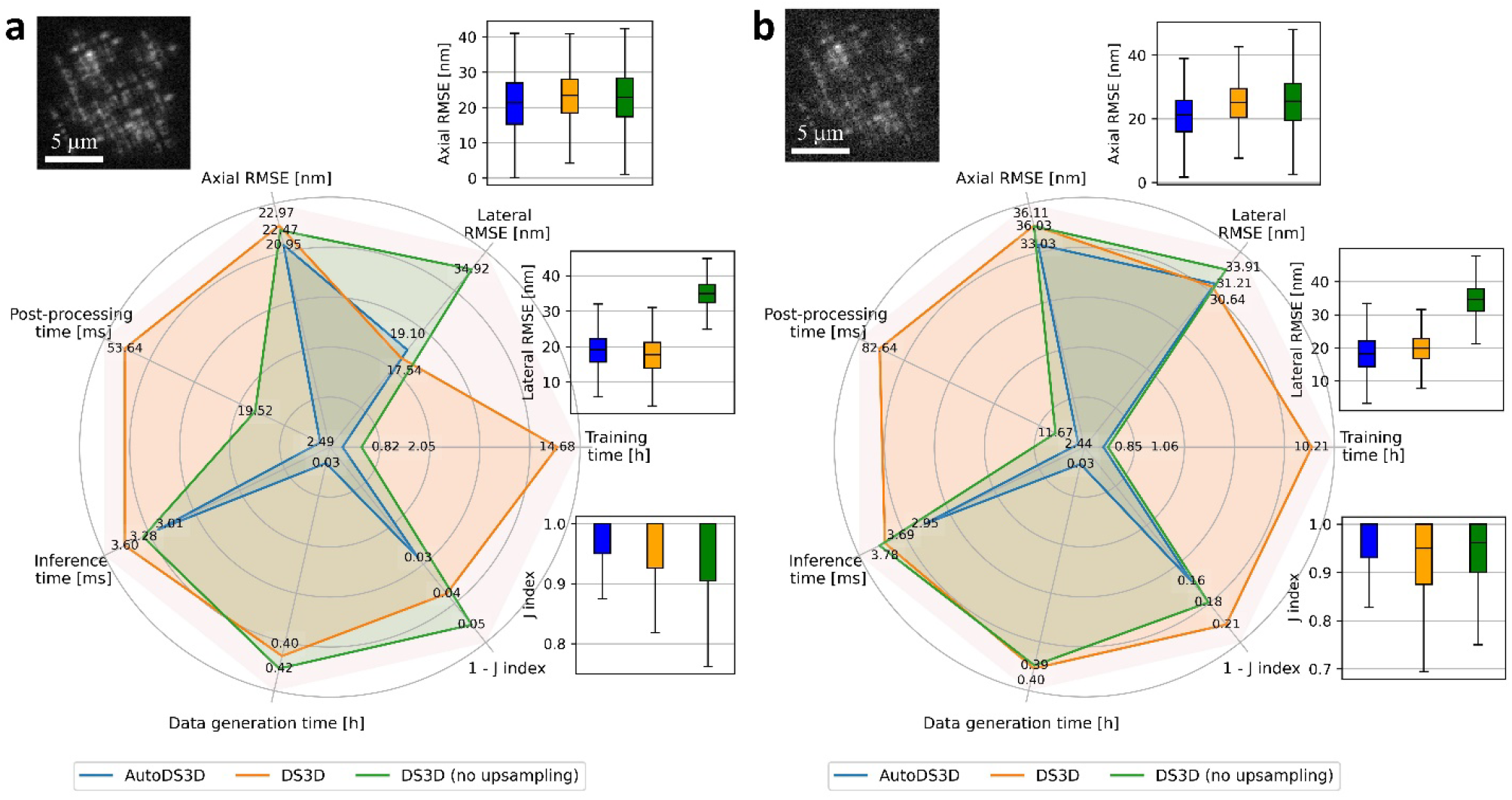
Comparison of accuracy and efficiency among AutoDS3D, DS3D and DS3D (no up-sampling) in simulation. Computation efficiency metrics include training data generation time, training time, inference time, post-processing time, and accuracy metrics, including axial and lateral RMSE, as well as 1-Jaccard index, are analyzed through 1000 times of inference whose statistical box plots are shown around the radar plot: **a** for high SNR and **b** for low SNR. Example frames are shown for both cases.

Next, we validated AutoDS3D using experimental SMLM data with various PSFs. We started with a Tetrapod dataset^11^ (20k frames, 333 by 313 pixels, video 1) and processed it using both AutoDS3D and DS3D (with 4x up-sampling). As shown in Figure 4, both methods show comparable reconstruction quality. DS3D requires approximately 14 hours for this reconstruction, plus additional hours/days depending on the user expertise for parameter selection and validation. In contrast, AutoDS3D completes the process in less than 2 hours with a single click.

**Figure 4:**
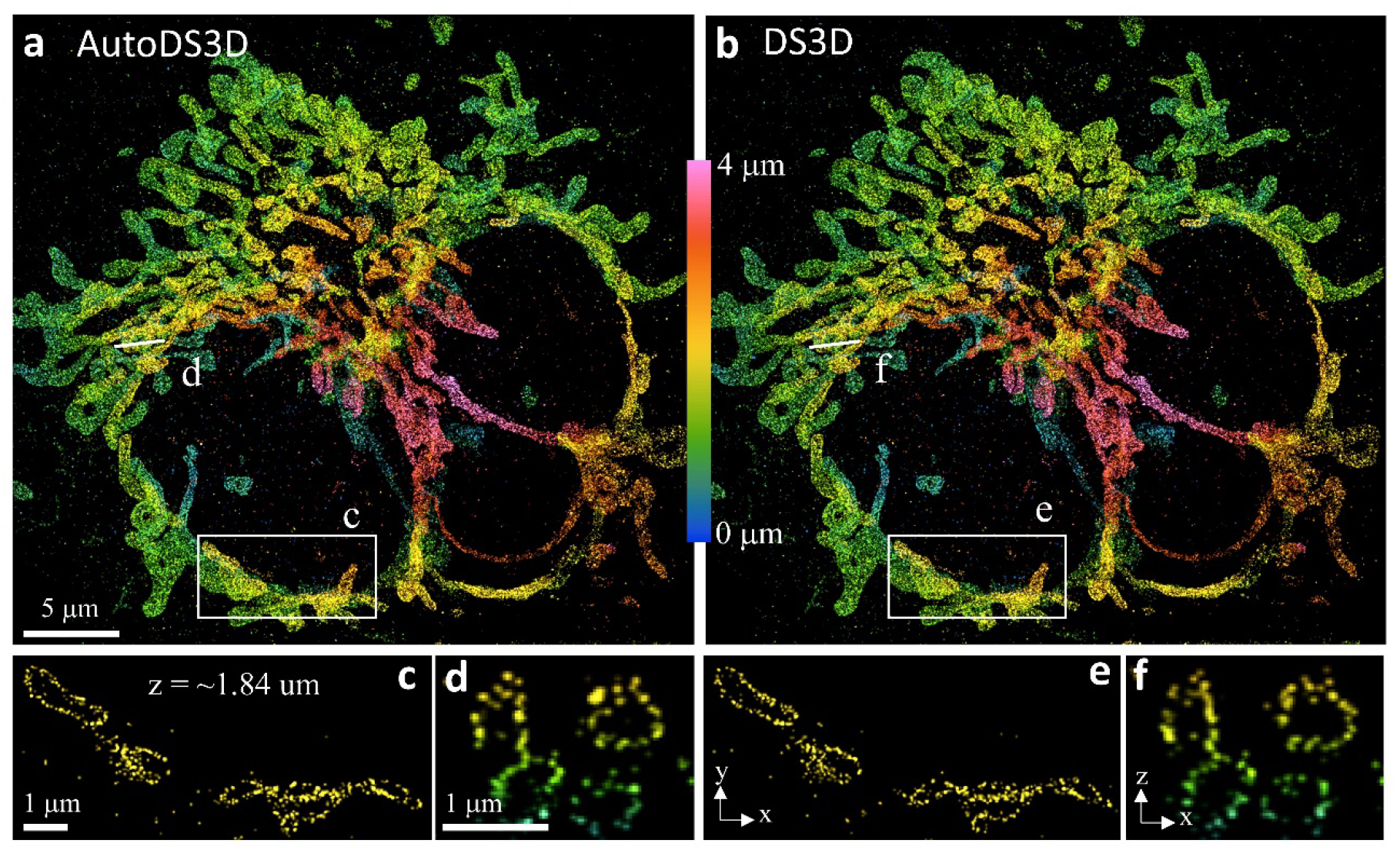
Experimental comparison between AutoDS3D and DS3D with tetrapod PSF. **a** reconstruction of AutoDS3D with cross section **d** and **c. b** reconstruction of DS3D with cross section **e** and **f** at the same location as **c** and **d**.

Next, we evaluated a dataset^27^ featuring a double-helix PSF, consisting of 80k frames (272 by 198 pixels, video 2). As shown in Figure 5, AutoDS3D matches DS3D in reconstruction quality while reducing processing time—completing the reconstruction in approximately 10 hours, compared to 50 hours for DS3D.

**Figure 5:**
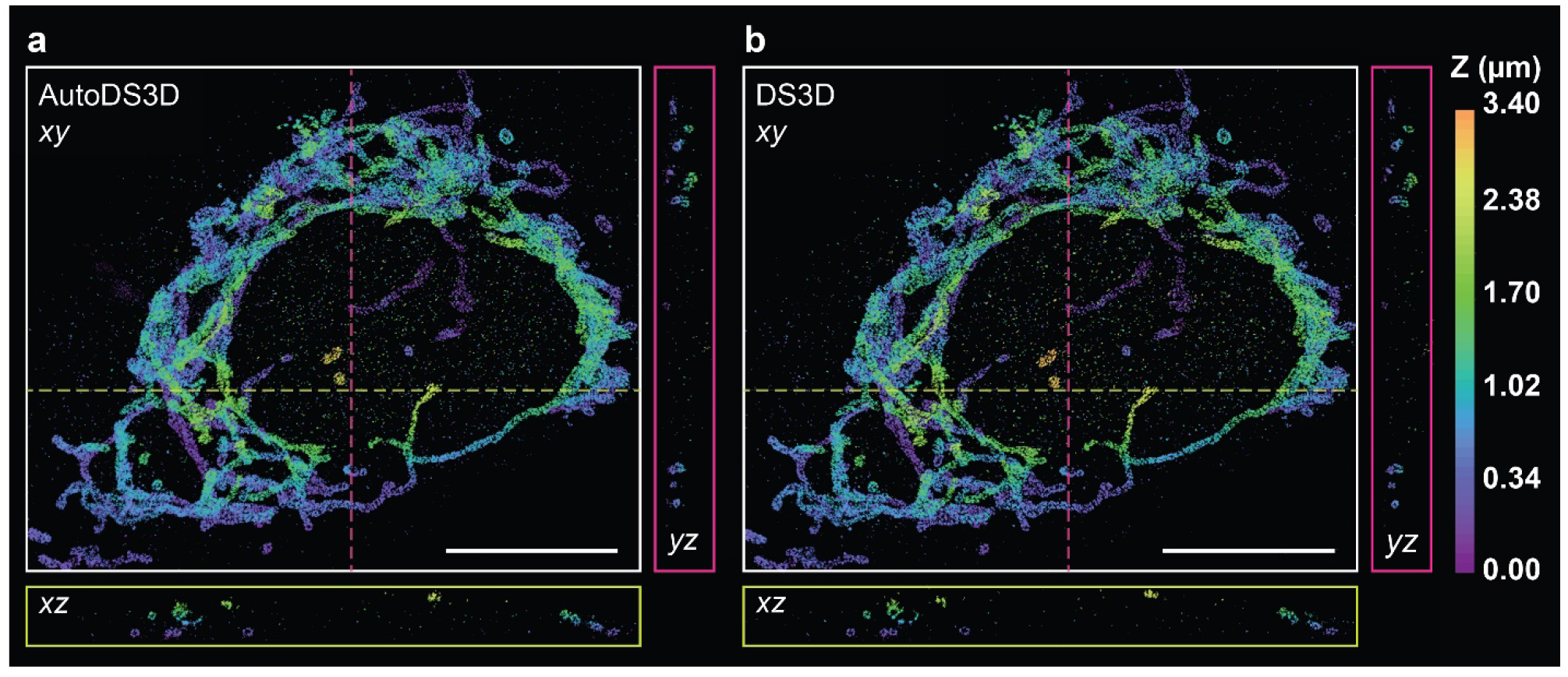
Experimental comparison between **a** AutoDS3D and **b** DS3D with double-helix PSF. The *xz* and *yz* views show 300 nm thick *xz* and *yz* cross sections along the dashed yellow and magenta lines shown in the xy view, respectively. Scale bars are 10 µm.

Finally, we evaluated AutoDS3D on two datasets featuring astigmatic PSFs^28^ (video 3 and video 4). In the reconstruction of nuclear pore complexes (Figure 6a), the two-layer structures with an axial separation of ∼50 nm are clearly resolved. The reconstructed microtubules (Figure 6b, with cross-sections in c and d) exhibit well-defined hollow tubular structures with a diameter of ∼50 nm.

**Figure 6:**
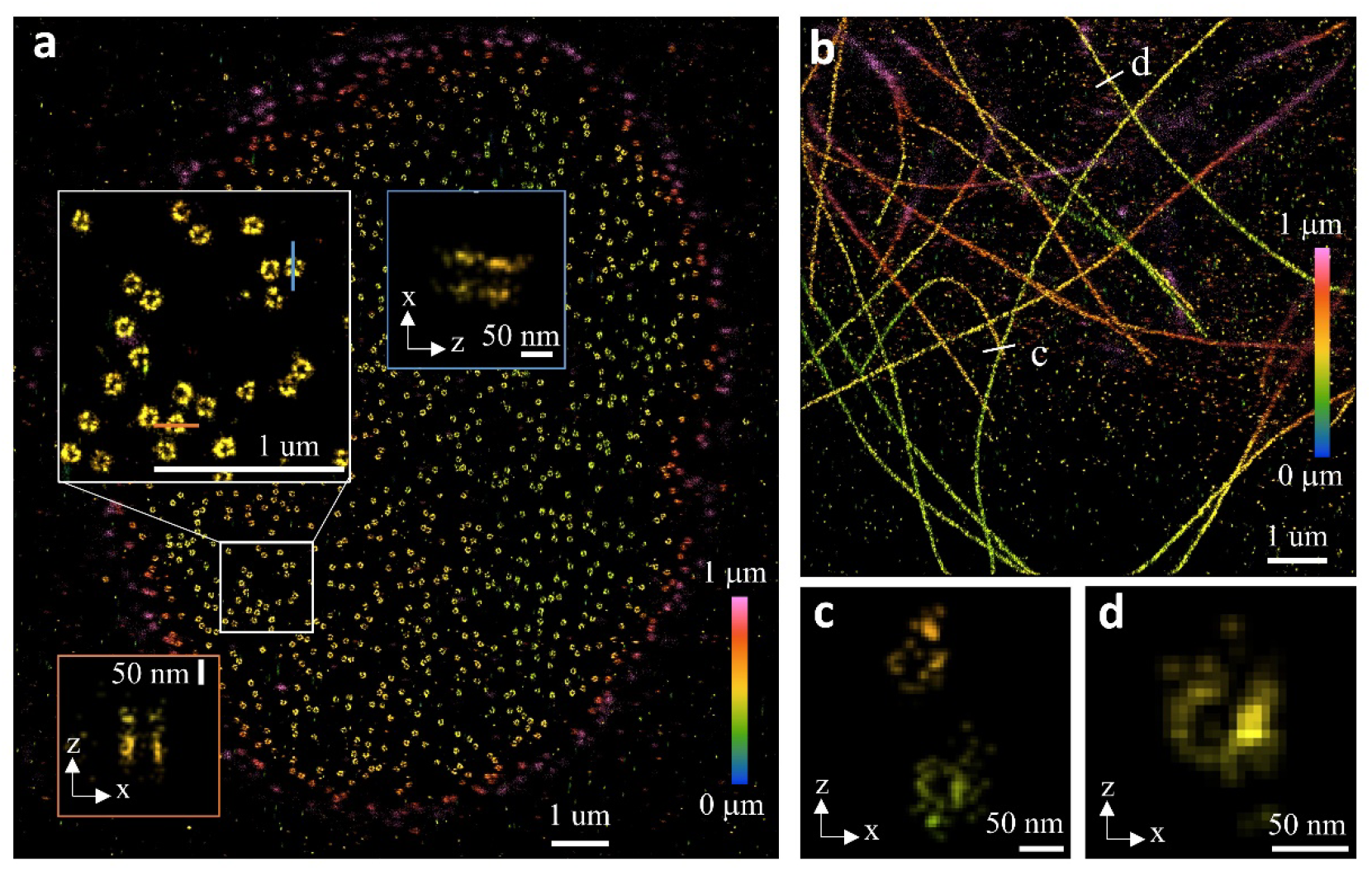
Experimental validation of AutoDS3D with an astigmatic PSF. **a** reconstruction of nuclear pore complexes with a zoom-in view and cross sections showing two-layer structures with axial distance of around 50 nm. **b** reconstruction of microtubules with axial cross section **c** and **d** showing hollow tubule structures.

## Discussion

We present several key contributions to the automation of deep-learning-based SMLM algorithms. The key results include: 1. complete removal of user involvement in 2D and 3D SMLM reconstruction, 2. alleviation of post-acquisition network training in the 2D case, and avoidance of dense sampling in the 3D case, leading to significantly faster reconstruction, and 3. patch-wise model selection in the 2D case to handle the locally heterogenous emitter density that is intrinsic to SMLM data of biological samples.

AutoDS is preloaded with four NN models for localization prediction in different experimental conditions. Users of AutoDS can use the set of pre-trained models offered in the package, while expert users may opt to train their own set of models and model selection logic to optimize their network performance for their needs. More generally, one can think of various multi-model methods that can be broadly applied to all manners of PSF-based SMLM datasets, regardless of biological structure, for better quality of image prediction.

Furthermore, the most time- and resource-intensive component of NNs is the preparation of training data and model training. Training data is the most important aspect of a well performing model; however, commonly, high quality ground truth data matching the experimental data is not easily available. Additionally, in Deep-STORM (both 2D and 3D), the generation of training data is human-supervised, namely, the user must observe the training data and visually assert that it resembles the experiment. By implementing pre-trained models in this workflow, we remove the requirement for model training, which eliminates the need for preparing training datasets and saves on computing and human resources.

Taken together, AutoDS provides better functionality, improved output quality, and user-friendliness compared to the single-model module. This multi-model, patch-based approach can also be easily extended to other image-based NNs that are confronted with heterogeneous image data from cell biology samples, such as in denoising^29,30^, image restoration^19^, and segmentation^31–33^. Future implementations of AutoDS can include an automated and non-rigid patch size selection module to confine similar structural densities within a single patch for more accurate model application.

In the 3D case, using a Gaussian representation of molecules, AutoDS3D resolves the trade-off between localization precision and computational efficiency, allowing the use of larger voxel sizes without compromising accuracy. AutoDS3D integrates the key steps of 3D SMLM image processing into a user-friendly GUI, making advanced localization accessible to users without computational expertise.

Although AutoDS3D enables one-click 3D super-resolution reconstruction, it still requires time-consuming model training. One future direction is to incorporate pre-trained models, similar to the 2D case, to enable instantaneous inference without the need for training. However, a key challenge in this approach will be ensuring compatibility between the PSF models used in the pre-trained models and those encountered in practical experiments on the user side.

## Methods

### Parameter extraction and model selection

Prior to patch parameter estimation, we evaluate some global experimental parameters, which are the means and standard deviations of: the noise, the emission intensity, and the PSF width (sigma), as well as a rough estimation of the overall emitter density in the video. These predicted parameters are later used for the classification of the emitter density and the signal-to-noise ratio (SNR) in each patch. To extract those parameters, we fit the first 100 emission events to the PSF model using a non-linear least squares algorithm. Then, we extract the PSF width (the standard deviation, assuming a 2D Gaussian model) by averaging the estimated value over 100 fits. Importantly, Gaussian fitting is a compute-expensive process; therefore, we only use it to extract relevant PSF features from a limited set of localizations. The emitters in each patch are located using much simpler methodology, namely, by searching local maxima in the patch.

The density-adaptive parameter extraction algorithm receives as input patches containing low-resolution single-molecule images and the extracted PSF features. First, we roughly estimate the number of emitters in each patch by looking for local maxima with intensity higher than the image (the entire field-of-view) mean intensity value plus two standard deviations. The emitter density is predicted by dividing the number of emitters by the analyzed patch size (as seen in Supplementary figure S1). Next, we resolve the intensity statistics of the emission event signal and the noise by creating a binary mask containing zeros in noise-related pixels and ones in signal-related pixels. The calculation of the mean and standard deviation of the noise is calculated using the noise-related pixels in each patch (zero valued pixels in the generated mask), and the signal mean and standard deviation is calculated using the signal-related pixels.

For the model selection, we focused on two extracted parameters: the emitter density and the SNR, defined by the mean signal amplitude divided by the noise standard deviation. While different networks could be trained for different combinations of these two parameters, we found that it was practically sufficient to train four AutoDS models with varying density and SNR conditions to analyze patches with different difficulty levels (see Table 2). We empirically chose the minimal SNR value to be 2 since the emitter signal is barely visible in this scenario; furthermore, we chose the maximal SNR value to be 8 since this SNR value is sufficient for precise emitter localization.

**Table 2:** The pre-trained model parameters. As the difficulty level increases, the emitter density increases, and the signal-to-noise ratio decreases. Decoupling those two parameters and producing a larger number of pre-trained nets did not make a substantial difference to the results.

|  | Model 1 | Model 2 | Model 3 | Model 4 |
| --- | --- | --- | --- | --- |
| Emitter density<br>Emitters $\left[\frac{1}{\mu m^2}\right]$ | 0.5 | 1.0 | 1.5 | 2.0 |
| Signal-to-noise ratio | 8 | 6 | 4 | 2 |

Each input patch is analyzed by the algorithm described above and is linked to a difficulty level between 1 and 4. Then, we average between the emitter density-based difficulty level and the SNR-based level to determine which pre-trained model will analyze the input patch.

### Image datasets

All 2D DNA-PAINT datasets were reused from a previous study^17^, in keeping with the principle of data reusability and reproducibility, as well as reducing laboratory materials and animal use. Animal handling from the previous work was performed according to the regulations by the Regierungspraesidium Karlsruhe. Data from the previous work is available on the Zenodo repository^34^. No animal studies were performed in this work. Methods ranging from sample handling to DNA-PAINT imaging outline the methods from previous study.

### Ground truth and predicted images

Ground truth α-tubulin and TOM20 images (low-emitter density, 0.5 nM imager strands, 10,000 frames; Figure 2) were fitted and reconstructed in Picasso software^22^. High-emitter density DNA-PAINT datasets for α-tubulin (5 nM imager strands, 400 frames) and TOM20 (10 nM imager strands, 400 frames) were input into the AutoDS NN in the Google Colab notebook environment (Pro version) for super-resolution image prediction. The “patchwise_analysis” parameter is selected which uses the set of pretrained models in the “Models” folder. The pixel size and emission wavelength of the dataset were defined as 233 nm, and the number of patches used was 8. For the single model comparisons, and Deep-STORM notebook was used to analyze the same high-emitter density datasets. The model upsampling factor of 8 determines the predicted image pixel size.

### Image analysis

Ground truth and predicted images of tubulin and TOM20 (Figure 2) were processed in Python. The image intensities were normalized, clipped in the range of the 0^th^ and the 99^th^ percentiles, and a Gaussian blur of sigma = 2 was applied. All images were registered using fiducial markers. Predicted images were compared against ground truth images.

### SNR characterization of AutoDS3D

In AutoDS3D, SNR characterization that is implemented after PSF characterization and background removal of raw images is crucial for training image generation. It involves noise parameters and photon count parameters. The noise is modeled by a Poisson noise and an additive Gaussian noise with mean and standard deviation. Because the SNR is determined by many factors, such as PSF model, brightness of molecules, PSF overlapping related to molecule density etc., we choose to control the molecule density and focus on a region of interest (ROI) with sparse molecules, ideally no PSF overlapping.

This ROI can be defined either by dragging a square in the pop-up image demonstration window or filling the parameter in the GUI. The ROI is cropped from the last 1000 frames of the image sequence and each pixel in the ROI forms a time sequence of length 1000. Then pixel-size mean and standard deviation are calculated and the pixel with the smallest mean value is seen as a noise pixel whose mean and standard deviation are used for the additive Gaussian noise. Empirically, we use a range (1.0, 1.4) times of the values and uniformly sample in that range. For photon count parameters, we first find the maximum pixel value (MPV) in the ROI. Then according to the retrieved PSF model, we calculate the linear factor between the photon count and the maximum pixel value of the PSF at the central z position of the defined z range. This linear factor is used to find the corresponding photon count of the experimental MPV. We also use a range of (0.5, 1.1) times of the detected photon count.

### The influence of nominal focal plane on the reconstruction in AutoDS3D

We examine the impact of the nominal focal plane (NFP) parameter on reconstruction by assigning different NFP values in the GUI. The analysis (Supplementary figure S5) shows that the z-range size remains relatively constant for consistent PSF shapes and the z-range position shifts linearly with NFP.

## Code availability

The software used in this work is publicly available at: https://github.com/alonsaguy/One-click-image-reconstruction-in-single-molecule-localization-microscopy-via-deep-learning.

## Acknowledgements

This research was supported by the Israel Science Foundation (grant No. 1081/24) and by partial financial support from the National Institute of General Medical Sciences of the National Institutes of Health (Grant R35GM155365) and startup funds from the Cancer Prevention and Research Institute of Texas (Grant RR200025) to A.-K.G. M.H. acknowledges funding by the Deutsche Forschungsgemeinschaft (DFG) (grants GRK 2566, CRC 1177). Y.S. is supported by the Zuckerman Foundation. A.S. is supported by the Fulbright Program under a Fulbright Visiting Scholar Program.

## References

1. Rust, M. J., Bates, M. & Zhuang, X. Sub-diffraction-limit imaging by stochastic optical reconstruction microscopy (STORM). Nature Methods 3, 793–795 (2006).

2. Heilemann, M. et al. Subdiffraction-Resolution Fluorescence Imaging with Conventional Fluorescent Probes. Angewandte Chemie International Edition 47, 6172–6176 (2008).

3. Jungmann, R. et al. Single-molecule kinetics and super-resolution microscopy by fluorescence imaging of transient binding on DNA origami. Nano Letters 10, (2010).

4. Betzig, E. et al. Imaging intracellular fluorescent proteins at nanometer resolution. Science 313, 1642–1645 (2006).

5. Hess, S. T., Girirajan, T. P. K. & Mason, M. D. Ultra-High Resolution Imaging by Fluorescence Photoactivation Localization Microscopy. Biophysical Journal 91, 4258–4272 (2006).

6. Sauer, M. & Heilemann, M. Single-Molecule Localization Microscopy in Eukaryotes. Chem Rev 117, 7478–7509 (2017).

7. Pavani, S. R. P. et al. Three-dimensional, single-molecule fluorescence imaging beyond the diffraction limit by using a double-helix point spread function. Proceedings of the National Academy of Sciences 106, 2995–2999 (2009).

8. Shechtman, Y., Sahl, S. J., Backer, A. S. & Moerner, W. E. Optimal point spread function design for 3D imaging. Physical Review Letters 113, (2014).

9. Nehme, E., Weiss, L. E., Michaeli, T. & Shechtman, Y. Deep-STORM: super-resolution single-molecule microscopy by deep learning. Optica 5, 458 (2018).

10. Speiser, A. et al. Deep learning enables fast and dense single-molecule localization with high accuracy. Nature methods 18, 1082–1090 (2021).

11. Nehme, E. et al. DeepSTORM3D: dense 3D localization microscopy and PSF design by deep learning. Nature Methods 17, 734–740 (2020).

12. Saguy, A. et al. DBlink: dynamic localization microscopy in super spatiotemporal resolution via deep learning. Nat Methods 20, 1939–1948 (2023).

13. Nogin, Y. et al. DeepOM: single-molecule optical genome mapping via deep learning. Bioinformatics 39, btad137 (2023).

14. Saliba, N., Gagliano, G. & Gustavsson, A.-K. Whole-cell multi-target single-molecule super-resolution imaging in 3D with microfluidics and a single-objective tilted light sheet. Nature Communications 15, 10187 (2024).

15. Ferdman, B. et al. VIPR: vectorial implementation of phase retrieval for fast and accurate microscopic pixel-wise pupil estimation. Optics Express 28, 10179 (2020).

16. Albertazzi, L. & Heilemann, M. When weak is strong: a plea for low affinity binders for optical microscopy. Angewandte Chemie International Edition e202303390 (2023).

17. Narayanasamy, K. K., Rahm, J. V., Tourani, S. & Heilemann, M. Fast DNA-PAINT imaging using a deep neural network. Nature Communications 13, 5047 (2022).

18. Jang, S. et al. Neural network-assisted single-molecule localization microscopy with a weak-affinity protein tag. Biophysical Reports 3, 100123 (2023).

19. Weigert, M. et al. Content-aware image restoration: pushing the limits of fluorescence microscopy. Nature Methods 15, 1090–1097 (2018).

20. Fu, S. et al. Field-dependent deep learning enables high-throughput whole-cell 3D super-resolution imaging. Nature Methods 20, 459–468 (2023).

21. Xiao, D. et al. Large-FOV 3D localization microscopy by spatially variant point spread function generation. Science Advances (2024).

22. Schnitzbauer, J., Strauss, M. T., Schlichthaerle, T., Schueder, F. & Jungmann, R. Super-resolution microscopy with DNA-PAINT. Nature Protocols 12, 1198–1228 (2017).

23. Wang, Z., Simoncelli, E. P. & Bovik, A. C. Multiscale structural similarity for image quality assessment. in The Thrity-Seventh Asilomar Conference on Signals, Systems & Computers, 2003 vol. 2 1398-1402 Vol.2 (2003).

24. Prieto, G., Chevalier, M. & Guibelalde, E. MS_SSIM Index as a Java plugin for ImageJ. (2014).

25. Narayanasamy, K. K. et al. Visualizing Synaptic Multi-Protein Patterns of Neuronal Tissue With DNA-Assisted Single-Molecule Localization Microscopy. Front Synaptic Neurosci 13, 671288 (2021).

26. Liu, S. et al. Universal inverse modeling of point spread functions for SMLM localization and microscope characterization. Nat Methods 21, 1082–1093 (2024).

27. Nakatani, Y., Gaumer, S., Shechtman, Y. & Gustavsson, A.-K. Long-Axial-Range Double-Helix Point Spread Functions for 3D Volumetric Super-Resolution Imaging. J. Phys. Chem. B 128, 11379–11388 (2024).

28. Li, Y. et al. Real-time 3D single-molecule localization using experimental point spread functions. Nature Methods 15, 367–369 (2018).

29. Krull, A., Buchholz, T.-O. & Jug, F. Noise2void-learning denoising from single noisy images. in 2129–2137 (2019).

30. Goncharova, A. S., Honigmann, A., Jug, F. & Krull, A. Improving Blind Spot Denoising for Diffraction-Limited Microscopy Data. in.

31. Falk, M. et al. Heterochromatin drives compartmentalization of inverted and conventional nuclei. Nature 570, 395–399 (2019).

32. Stringer, C., Wang, T., Michaelos, M. & Pachitariu, M. Cellpose: a generalist algorithm for cellular segmentation. Nature Methods 18, 100–106 (2021).

33. Zakrzewski, F. et al. Automated detection of the HER2 gene amplification status in Fluorescence in situ hybridization images for the diagnostics of cancer tissues. Scientific Reports 9, 8231 (2019).

34. K. K. Narayansamy. Accelerating DNA-PAINT imaging with a deep neural network. 10.5281/zenodo.6966132.

