## Supplementary information for "One-click image reconstruction in single-molecule localization microscopy via deep learning"

**Supporting information**

In this work, we aimed to enhance the existing implementation of Deep-STORM^1^ by using a patch-wise processing scheme, which divides whole image frames into patches for prediction; however, the patch-wise implementation came with new challenges. For example, localization algorithms suffer from reduced performance at the edges of the analyzed patches because the PSF is partially truncated by the image edge, rendering the truncated PSF difficult to fit. This problem can be neglected in full FOV analysis since inaccuracies appear only at the edges of the image. However, in our case this problem had a greater impact on reconstruction quality and had to be addressed. Therefore, we employed patch overlaps and ignored localization at the edges of each patch. Specifically, we used a 4-pixel overlap between neighboring patches and ignored the network predictions within 2 pixels from the patch edges, thereby producing a final image with a continuous structure.

Another challenge that arises when analyzing small patches instead of images is the frequent occurrences of patches that do not contain any emitters. Deep-STORM pre-processing pipeline involves rescaling the pixel values to the [0,1] range, and matching the mean and standard deviation to the empirical values in the training data. Unfortunately, the absence of emitters in a patch with a relatively high noise variance leads to the hallucination of emitters by Deep-STORM. To overcome this challenge, we employed a minimum subtraction filter for each pixel in the field-of-view; furthermore, at the model selection stage, we normalized the patches according to the entire field-of-view intensities and assigned probable “noise-only” patches to the model that was trained on high-SNR scenario. This led to “conservative” prediction of emitter locations and significant reduction of hallucinated emitters in the reconstruction.

In addition, the implementation of the patch-wise analysis in Deep-STORM required modifications in the upsampling layer of the network architecture. Deep-STORM’s model was initially trained to convert upsampled low-resolution blinking frames to upsampled high-resolution localization heatmaps. To obtain upsampled low-resolution frames, Deep-STORM’s pipeline used Nearest Neighbor upsampling method; hence, creating non-realistic observations. We modified Deep-STORM’s model architecture to alleviate the need of this step; therefore, instead of using a fully symmetric encoder-decoder architecture, we added an adjustable number of upsampling layers after the last decoder layer to enable the model to learn an upsampling function by itself. The number of upsampling layers is determined by the selected scale factor.

Finally, since Deep-STORM’s initial release, most of the Python packages and relevant hardware have drastically changed. Therefore, we have updated the implementation of some parts of Deep-STORM to accommodate modern package versions and hardware. Apart from the changes mentioned above we did not alter the logic of Deep-STORM’s pipeline, i.e., most of our updates were meant for improving Deep-STORM’s compatibility with modern Python packages.

**Supplementary figures and tables**

|  | Mean noise | Noise STD | Mean signal | Signal STD | Emitter density |
| --- | --- | --- | --- | --- | --- |
| $\alpha$-tubulin | 781.5 | 645.1 | 4387.3 | 988.4 | 0.25 |
| TOM20 | 403.2 | 146.0 | 1700.4 | 549.1 | 0.20 |

Table S1: Experimental parameter estimation extracted by our method for the two DNA-PAINT experiments reported in Figure 2. We extract the mean/standard deviation of the intensity of noise related pixels, the mean/standard deviation of the intensity of signal related pixels, the emitter density.

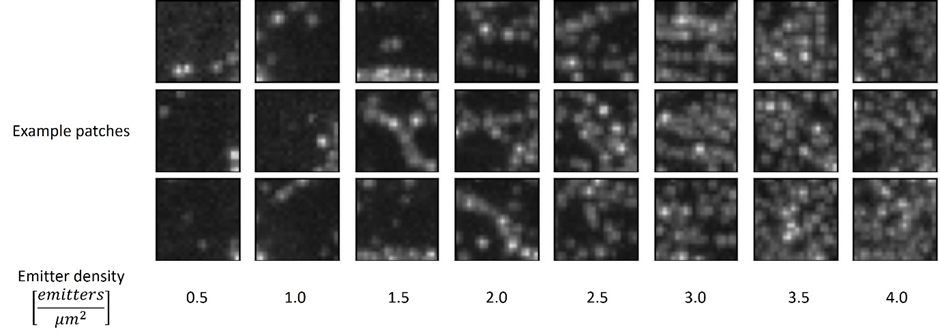

Figure S1: Visualization of representative emitter density classifications made by AutoDS pipeline. We used sparse SMLM experimental data to generate a semi-simulated SMLM video with high emitter density by summing every 120 frames. Then, we used AutoDS pipeline to estimate the emitter density over different patches. Each column shows three representative patches that matched the estimated emitter density indicated at the bottom. The patch size is 2.5x2.5 μm^2^.

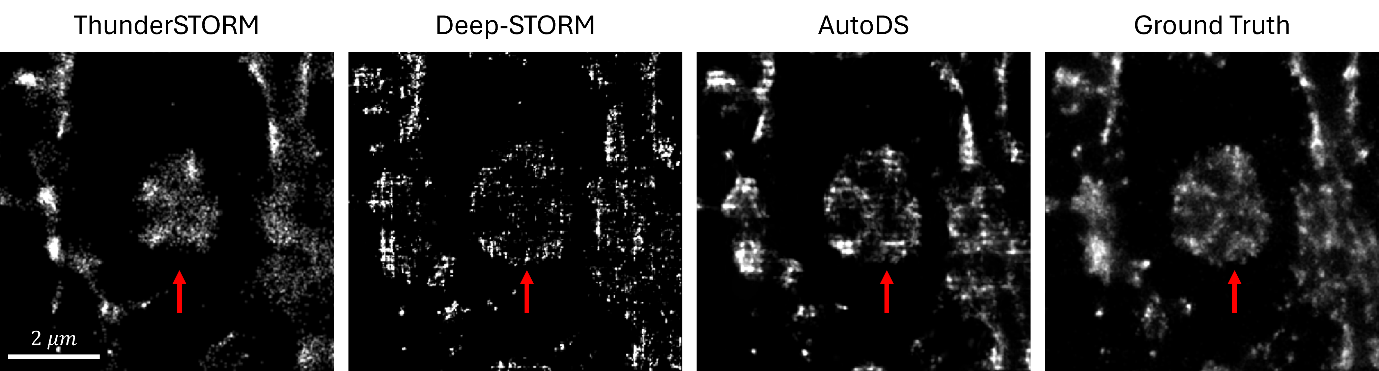

Figure S2: Demonstration of AutoDS reconstruction superiority in comparison to Deep-STORM and ThunderSTORM^2^. Left to right: ThunderSTORM, Deep-STORM, and AutoDS reconstructions of a high-density sample with labeled $\alpha$-tubulin, and the ThunderSTORM reconstruction of the same field of view in low emitter density scenario. Red arrow marks a region where ThunderSTORM fails to capture the tubulin structure, probably due to its incompatibility with higher densities; although Deep-STORM manages to partially reconstruct a structure like the ground truth, AutoDS reconstruction has much higher fidelity to the ground truth.

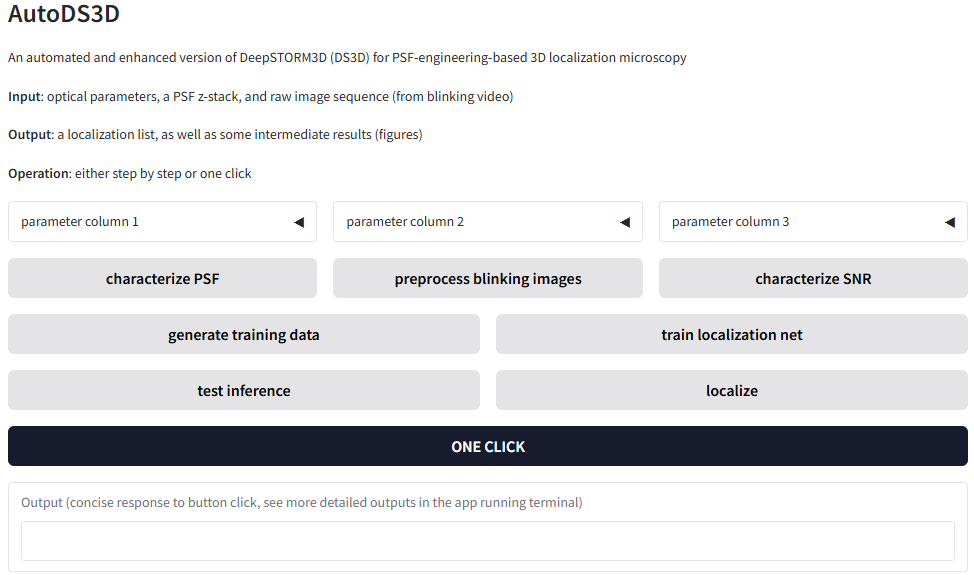

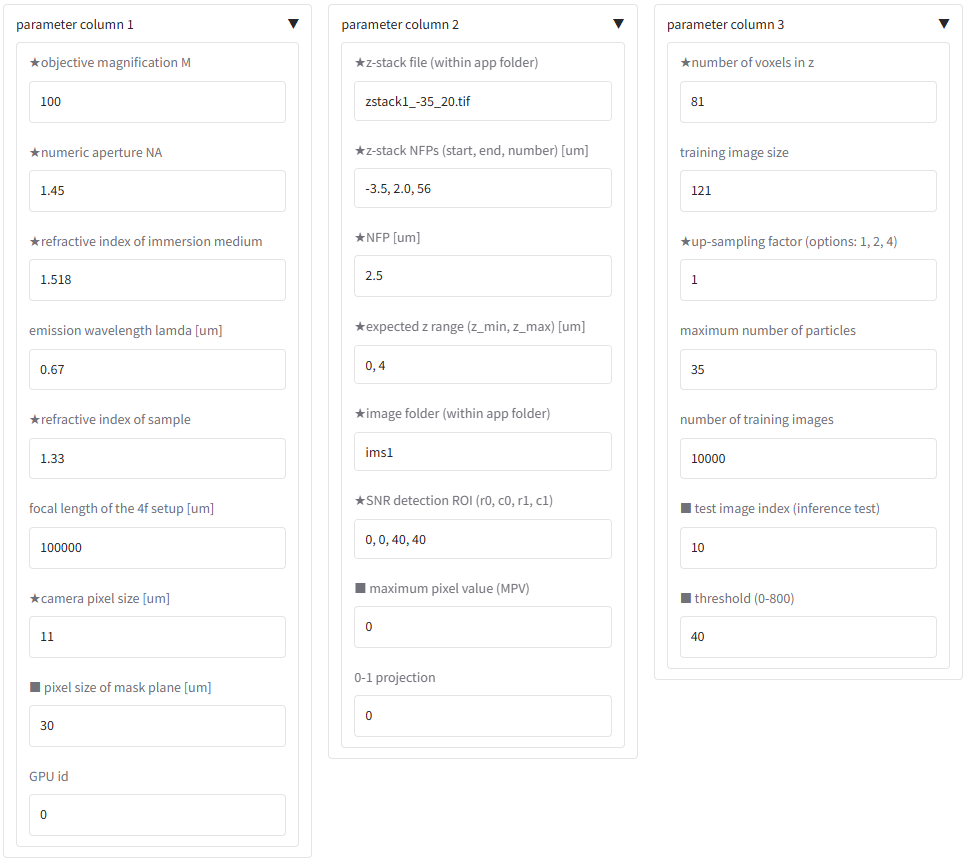

Figure S3: Graphic user interface (GUI) of AutoDS3D and the unfolding parameter columns.

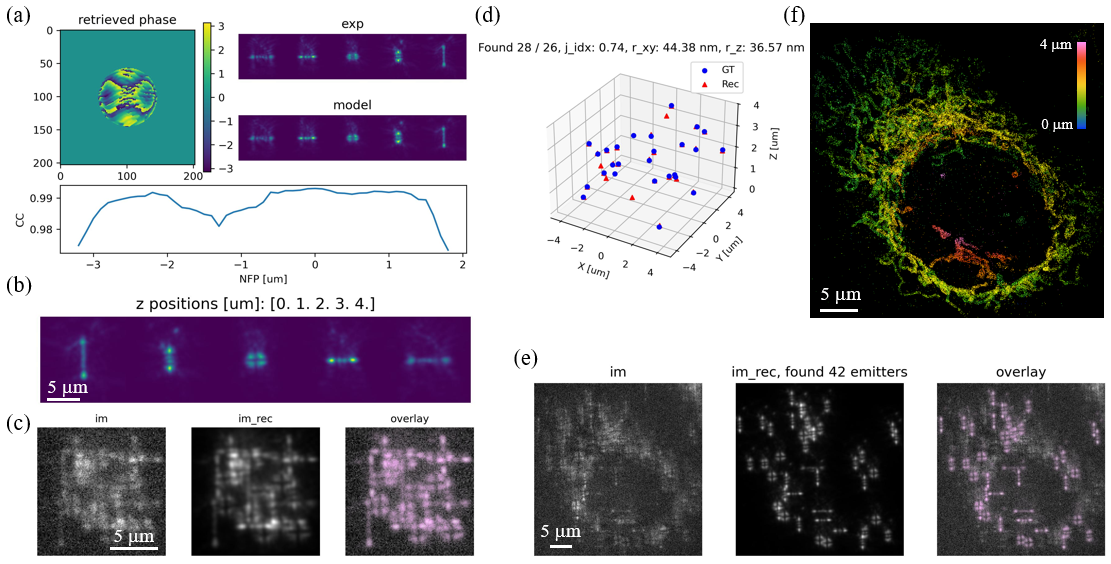

Figure S4: Intermediate results when running AutoDS3D to process a dataset with low signal to noise ratio. **a**, phase retrieval result where CC denotes correlation coefficient. **b**, PSFs across the z range. **c-d**, inference and reconstruction of the trained model on a simulated image. **e**, reconstruction of an experimental image. **f**, 3D reconstruction of the sample structure.

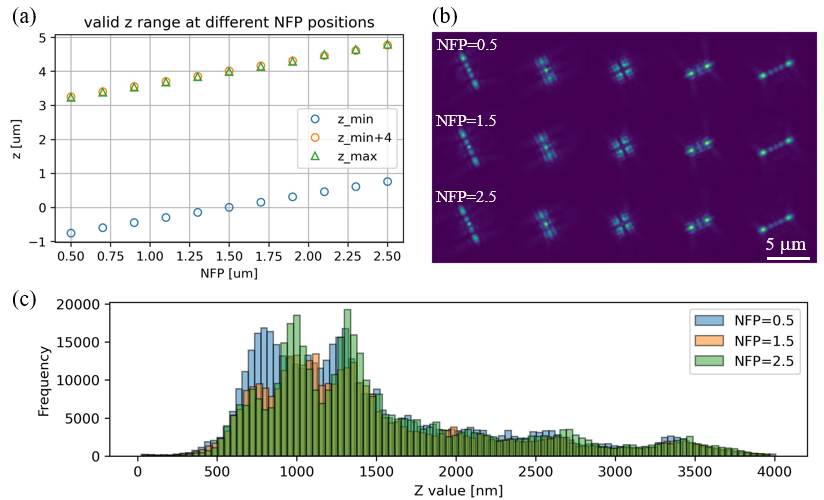

Figure S5: Analysis of the influence of nominal focal plane (NFP) on reconstruction. NFP=1.5 is the reference position and its PSFs in the z range of (0, 4) µm are the reference PSF shapes. For each NFP value deviating from the reference position, we detect the z range corresponding to the reference PSF shapes, as shown in **a** and **b**. For NFP of 0.5, 1.5, and 2.5 µm, we train the network accordingly and test the trained network on the data in Figure 4. The z histograms of the localizations **c** show similar features.

**References**

1. Nehme, E., Weiss, L. E., Michaeli, T. & Shechtman, Y. Deep-STORM: super-resolution single-molecule microscopy by deep learning. *Optica* **5**, 458–464 (2018).

2. Ovesný, M., Křížek, P., Borkovec, J., Švindrych, Z. & Hagen, G. M. ThunderSTORM: a comprehensive ImageJ plug-in for PALM and STORM data analysis and super-resolution imaging. *Bioinformatics* **30**, 2389–2390 (2014).
